# Common genetic variation drives molecular heterogeneity in human IPSCs

**DOI:** 10.1101/055160

**Authors:** Helena Kilpinen, Angela Goncalves, Andreas Leha, Vackar Afzal, Sofie Ashford, Sendu Bala, Dalila Bensaddek, Francesco Paolo Casale, Oliver Culley, Petr Danacek, Adam Faulconbridge, Peter Harrison, Davis McCarthy, Shane A McCarthy, Ruta Meleckyte, Yasin Memari, Nathalie Moens, Filipa Soares, Ian Streeter, Chukwuma A Agu, Alex Alderton, Rachel Nelson, Sarah Harper, Minal Patel, Laura Clarke, Reena Halai, Christopher M Kirton, Anja Kolb-Kokocinski, Philip Beales, Ewan Birney, Davide Danovi, Angus I Lamond, Willem H Ouwehand, Ludovic Vallier, Fiona M Watt, Richard Durbin, Oliver Stegle, Daniel J Gaffney

## Abstract

Induced pluripotent stem cell (iPSC) technology has enormous potential to provide
improved cellular models of human disease. However, variable genetic and phenotypic
characterisation of many existing iPSC lines limits their potential use for research and therapy. Here, we describe the systematic generation, genotyping and phenotyping of 522 open access human iPSCs derived from 189 healthy male and female individuals as part of the Human Induced Pluripotent Stem Cells Initiative (HipSci:http://www.hipsci.org). Our study provides a comprehensive picture of the major sources of genetic and phenotypic variation in iPSCs and establishes their suitability for use in genetic studies of complex human traits and cancer. Using a combination of genomewide analyses we find that 5-25% of the variation in different iPSC phenotypes, including differentiation capacity and cellular morphology, arises from differences between individuals. We also assess the phenotypic effects of rare, genomic copy number mutations that are recurrently seen following iPSC reprogramming and present an initial map of common regulatory variants affecting the transcriptome of pluripotent cells in humans.

## Introduction

Induced pluripotent stem cells (iPSCs) are important model systems for human disease ^1^. A critical unanswered question is whether iPSCs can be used to study the functions of genetic variants associated with complex disease and normal human phenotypic variation. Previous work has suggested that individual iPSC lines may be highly heterogeneous ^2–   5^. Substantial iPSC heterogeneity means that the subtle effects of common genetic variants might be hard to detect. Existing iPSC lines often have limited genetic and phenotypic data of variable quality, or are derived from individuals with severe genetic disorders, limiting their utility for studying other phenotypes. Although previous large-scale studies in pluripotent stem cells have been undertaken, they have not systematically derived hiPSCs at scale nor focused on characterising phenotypic effects of naturally occurring genetic variation ^5, 6^. Thus, there is a critical need for large, well-characterised collections of human iPSCs (hiPSCs) systematically generated using a single experimental pipeline.

To overcome this problem there is a requirement for large, well-characterised collections of human iPSCs (hiPSCs) that have been systematically generated using a single experimental pipeline. Furthermore, The Human Induced Pluripotent Stem Cells Initiative (HipSci; www.hipsci.org) was established to generate a large, high-quality, open-access reference panel of human iPSC lines. A major focus of the program is the systematic derivation of iPSCs from hundreds of healthy volunteers using a standardised and well-defined experimental pipeline. Each generated line is extensively characterised and lines with accompanying genetic and phenotypic data are available for use by the wider research community. Here, we report initial results from the characterization of the first 522 iPSC lines derived from 189 healthy individuals. Our study shows that common genetic variants produce readily detectable effects in iPSCs, and provides the first map of regulatory variation in human pluripotent stem cells. We also demonstrate that differences between donor individuals have pervasive effects at all phenotypic levels in iPSCs, from the epigenome, transcriptome and proteome to cell differentiation and morphology.

## Results

### Sample collection and IPSC derivation

Samples for the project were collected over a period of 13 months between February 2013 and March 2014 during which we received a total of 430 skin punch biopsies from healthy, unrelated research volunteers, the vast majority of which were of Northern European ancestry (**Extended Data Fig. 1**) recruited through the NIHR Cambridge BioResource(http://www.cambridgebioresource.org.uk). Fibroblast outgrowths from skin explants of each individual were reprogrammed using a Sendai viral vector system ^7^ on a feeder layer of mouse embryonic fibroblasts and 234 (54.4%) produced pluripotent colonies within 35 days post transduction on average. Unsuccessful reprogramming attempts were due to failure to produce fibroblast outgrowths (50 individuals, 11.6%) or failure to produce pluripotent colonies (146 individuals, 34.0%). Of the 234 successfully reprogrammed samples, 189 were sufficiently advanced in our experimental pipeline to be included in the current study.

We established multiple independent lines from most donors (92% of donors had > 1 line, 72% had 3 lines) resulting in a total of 522 iPSC lines that were subjected to an initial set of genetic and phenotypic assays (hereafter ‘Tier 1’ assays) (**Fig. 1a**). Tier 1 assays included array based genotyping and gene expression profiling of the iPSCs and their fibroblast progenitors. For 301 lines we quantified protein expression of NANOG, OCT4 andSOX2 using immunohistochemistry followed by quantitative image analysis using the Cellomics (Thermo Fisher Scientific) high content imaging system. We also differentiated 372 lines into neuroectoderm (dEC), mesoderm (dME), and endoderm (dEN), using a defined culture system ^8^, and measured the expression of three lineage-specific differentiation markers (**Fig. 1a**) using the Cellomics platform (**Extended Data Figure 2, Methods**).

The Tier 1 assay data were used to select 1-2 high quality lines for each donor for further phenotyping and cell line banking, minimising the number of genetic abnormalities and maximizing pluripotency. For this study, 167 lines (hereafter ‘selectedlines’) from 127 donors were selected based on Tier 1 assay data, and profiled using RNA-seq, with lines from 27 donors subjected to DNA methylation profiling, 9 donors to quantitative proteomics and 12 to cell morphological imaging (hereafter ‘Tier 2’ assays) (**Extended Data Figure 3, Supplementary Table 1, 2**).

**Figure 1.**
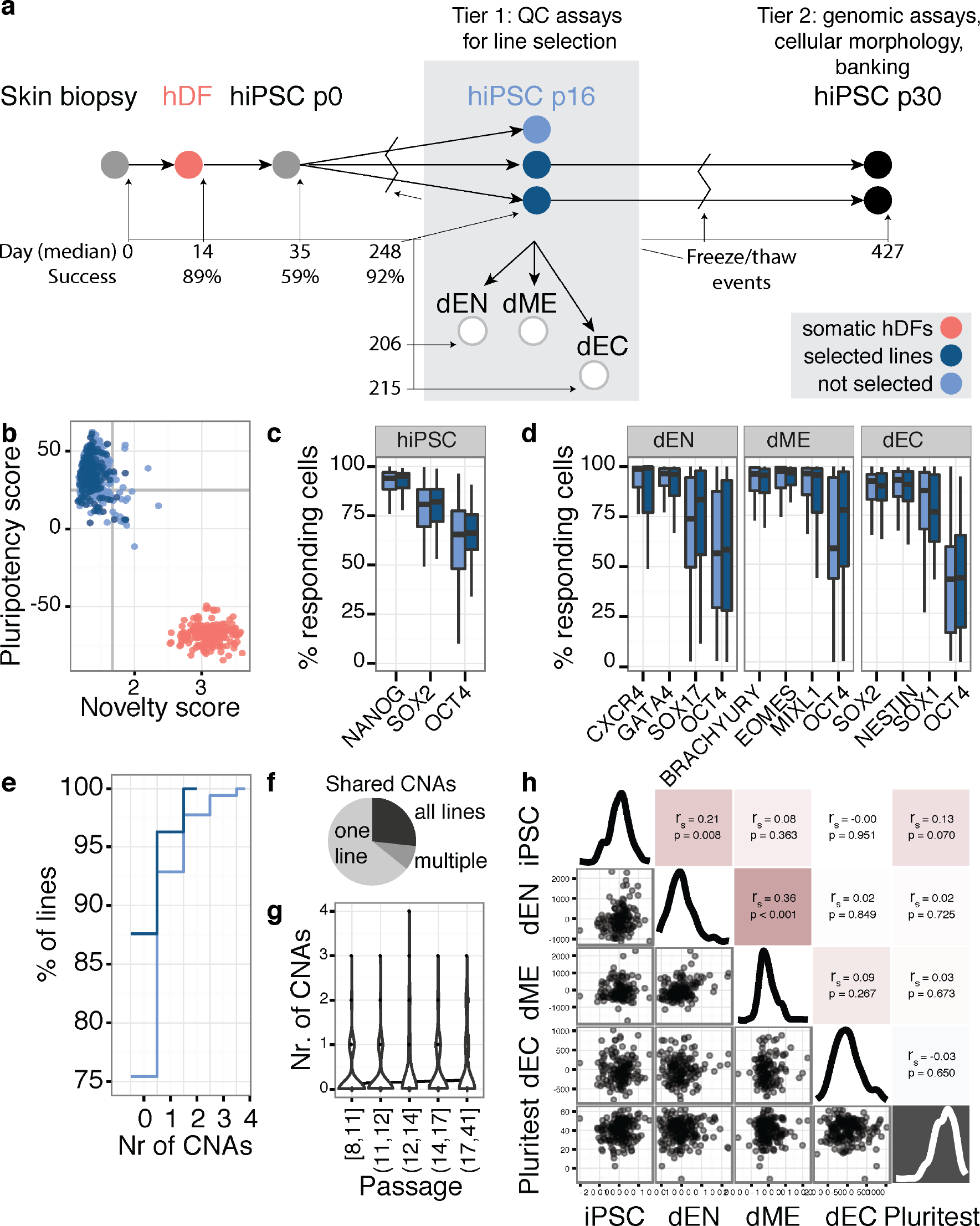
Experimental design of iPSC line generation and quality control. **(a)** Schematic of iPSC generation pipeline. hDF: human dermal fibroblasts; dEN: differentiated endoderm; dME: differentiated mesoderm; dEC: differentiated neuroectoderm. Samples for molecular profiling were taken at two stages: ‘Tier 1’ assays were profiled in cells at average passage 16, ‘Tier 2’ as sayswere carriedout at average passage 30 for selected lines (**Methods**). Shown below the x-axis is the day number (where receipt of a skin punch was day 0) for an average line to go from registration to each pipeline stage and corresponding success rates. Times do not reflect continuous periods in culture and include intervals where lines were frozen. Time to Tier 1 assay stage was defined as the mean day number when gene expression was profiled using microarrays. (**b-h**) Analyses of the Tier 1 quality control assays. **(b)** Pluripotency of lines assessed using PluriTest ^9^, a computational assay based on gene expression arrays. Comparison of PluriTest novelty score versus pluripotency score for 522 lines generated (light blue), for selected (dark blue) and hDFs (red). **(c)** Estimated fraction of iPSCs expressing pluripotency markers (% of responding cells) measured using immunostaining and high content imaging (**Methods**). **(d)** The percentage of responding cells stained for differentiation markers for endoderm (dEN),mesoderm (dME) and neuroectoderm (dEC). (**e-g**) Genetic stability in hiPSCs. **(e)** Distribution of number of CNAs across all lines (dark blue-selected lines, lightblue-lines not selected). **(f)** Fraction of CNAs shared by one, multiple or all clonal lines from the same donor. **(g)** Relationship between CNA count and line passage number. **(h)** Pairwise correlation between scores derived from immunostaining for pluripotency and differentiation and the PluriTest score.

### Pluripotency and genetic stability

We analysed Tier 1 gene expression array data using PluriTest ^9^, which suggested that all 522 lines displayed expression patterns typical of pluripotent cells (**Fig. 1b**). Using the Cellomics imaging data we quantified the fraction of cells expressing each marker and estimated that, on average, between 18% and 62% of cells in the iPSC lines expressed all three pluripotency markers NANOG, OCT4 and SOX2 (**Fig. 1c, S11**). The vast majority of lines (> 99%) successfully produced cells from all three germ layers during directed differentiation with the average line producing up to 70%, 84% and 77% of cells expressing all three markers of dEN, dME and dEC, respectively (**Fig. 1d**). Lineage-specific marker expression was positively correlated between endoderm and mesoderm as well as between endoderm markers and expression of NANOG, OCT4 and SOX2 in the original iPSCs (**Fig. 1h**). Together, these data indicate that virtually all of the hiPSC lines we have derived are pluripotent.

Aneuploidy and sub-chromosomal aberrations have frequently been observed in cultured pluripotent stem cells ^6, 10– 12^. We used genotyping arrays to detect copy number alterations (CNAs) between the 189 original fibroblasts and the 522 iPSCs lines derived from them, using a computational approach developed for this purpose ^13^. We estimate that we can detect genetic abnormalities of > 1Mb that occur in 20% or more cells within an individual line. Using this approach we called a total of 147 larger CNAs (> 1Mb in size). 4% of all lines (none of the lines selected for Tier 2 assays) had trisomies and 21% of all lines (12% of the selected lines) harboured one or more CNAs of, on average, 8.64Mb in length with duplications outnumbering deletions by 2.9 to 1 (**Fig. 1e**). Although the majority of CNAs are unique to single iPSC line, 36% are also observed in at least one replicate line from the same donor (with sharing defined as overlapping by at least one base), with 27% seen in all replicates (**Fig. 1f**). We found no significant association between the number of CNAs and any of passage number, donor age, gender and PluriTest score of a line (**Fig. 1g, Extended Data Fig. 4**).

### Recurrent copy number alterations in human IPSCs

CNAs observed in pluripotent stem cells (PSCs) are known to recur in specific genomic locations ^6, 11, 12^. Using our CNA call set, we next identified regions of recurrent genomic alteration. Our analysis builds on previous work in three ways. First, we obtained reference material from the donor, which was not collected in previous large-scale studies, enabling us to distinguish CNAs that have appeared during somatic development or reprogramming from germline variants.Second, our sample size (522 lines) was 14-fold larger than the largest reported sample of hiPSCs (37 hiPSCs in ref ^11^), four times larger than previous studies of pluripotent stem cells (PSCs) (136 PSC lines in ref ^6^), and twice as large as the biggest karyotyping study to date ^14^. Third, because we collected gene expression data in the same cells, we were able to characterize the downstream consequences of each recurrent CNA on gene expression.

We observed a number of regions where CNAs occurred significantly more often than expected under a uniform genomic distribution (**Methods**), including whole chromosome duplication of the × chromosome (six independent donors, p = 6.9×10^−6^), six sub-chromosomal duplications-on chr20q11.21 (13 donors, p = 1×10^−8^), chr17q (5 donors, p = 3.4×10^−5^), chr10q (4 donors, p = 1.2×10^−4^), chr1q32.1 (4 donors, p = 1.6×10^−5^), chr1q42.2 (3 donors, p = 5×10^−4^) and chr3q26 (3 donors, p = 8.9×10^−4^); and two regions of recurrent deletion at chr1q23.3 (2 donors, p = 4.9×10^−4^) and chr9q21 (2 donors, p = 1.2×10^−3^) (**Fig. 2a, Supplementary Table 3**). The six recurrent subchromosomal regions were between 0.8 and 6 Mb in length, with one comprising the short arm of chromosome 10 and another 84.5% of the long arm of chromosome 17. A number of the recurrent CNAs we detected have been previously observed in PSCs, including × trisomy ^6, 14, 15^ duplications of the long arm of chromosome 17 and a minimum amplicon on chromosome 20 ^6^.

**Figure 2.**
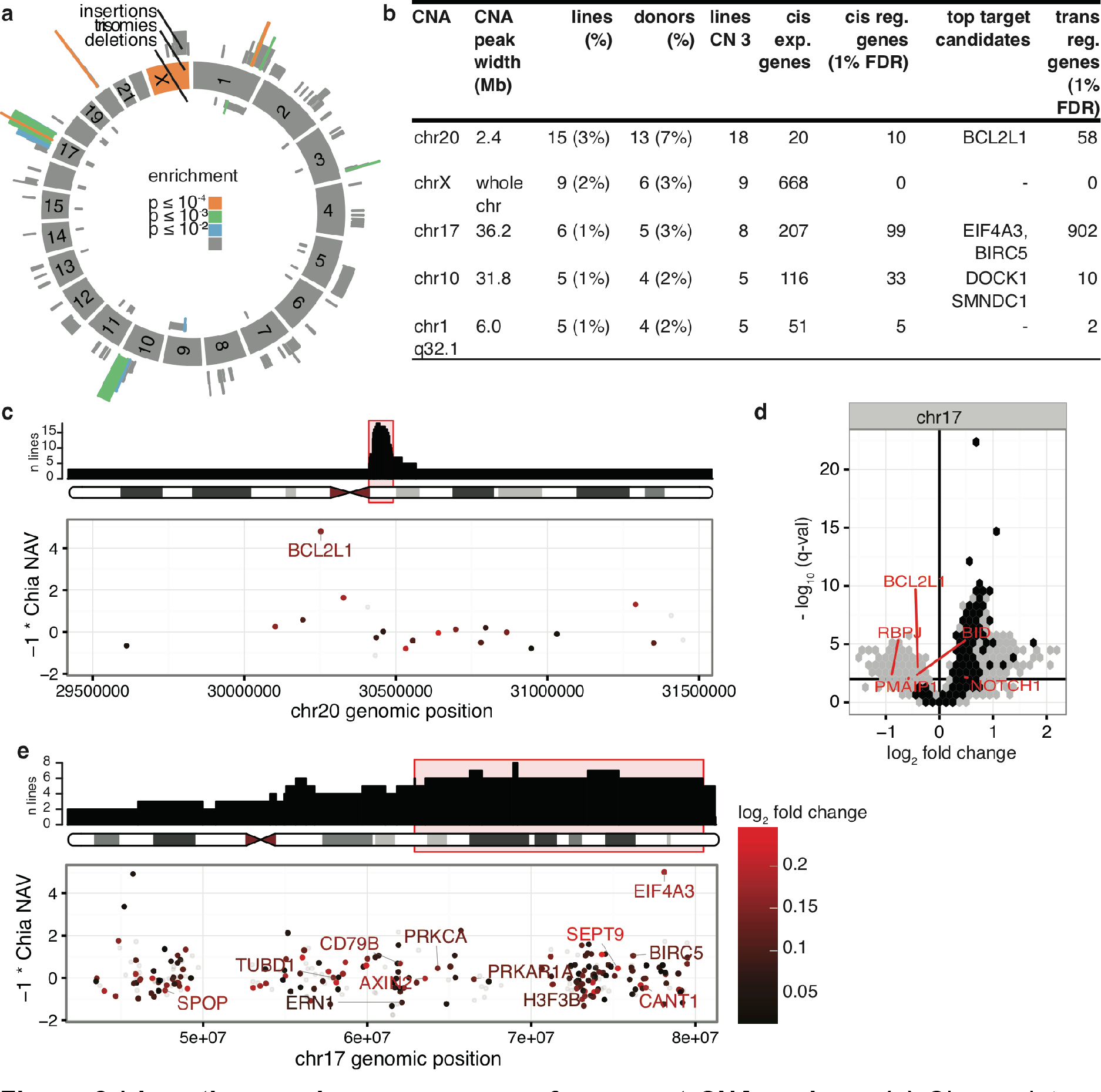
Locations and consequences of recurrent CNA regions. **(a)** Circos plot showing the genomic location of structural genetic alterations (copy number variants and trisomies) identified between each hiPSC line and the corresponding hDFs from which they were derived. Colours denote the significance level of recurrence of chromosomal (chromosome ring) and sub-chromosomal CNAs (frequency on the outside of the ring). **(b)** Selected recurrent CNAs and potential selection targetgenes in the region. **(c,e)** Location of the peak regions within the CNAs (top) and the genes expressed within these peak regions (bottom). In the top plot the y-axis denotes the number of lines with CN 3 (not necessarily a CNA as some donors have CN 3 on both the somatic cells and iPSCs). On the bottom plot the y-axis denotes the reduction in number of nuclei upon knockdown of a gene in a siRNA screen as a proxy phenotype for impact on cell proliferation. Highlighted are genes up-regulated when copy number increases and that are either known onco/tumour-suppressor genes or genes that score in the top 2% in ref.^16^. The colour code shows log_2_ gene expression fold change between the iPSC lines with copy number 2 and 3. **(d)** Differential expression of genes between lines with copy number 2 and 3 for the recurrent CNA on chromosome 17. Grey dots denote gene outside the CNA (regulated *in trans*), black dots denote genes inside the CNA (*in cis*). The significance level used (q = 0.01) is shown as a horizontal bar intersecting the y-axis.

Although recurrent CNAs could be due to mutational hotspots, we did not find significant overlap between our recurrent CNA set and annotated chromosome fragile sites ^17^ (p = 0.939, **Methods**). Recurrent CNAs could also arise if duplication or deletion of specific genes led to a selective advantage. For example, the chromosome 20 duplication is hypothesised to arise due to a growth advantage conferred by the overexpression of *BCL2L1*, a regulator of apoptosis^6, 18^. Consistent with this idea we found that 56% of CNA hotspots overlapped recurrent somatic copy number alterations in cancer, significantly more often than randomly generated control sets of equivalent size (18% overlap, p = 0.0095, lenient set from ^19^).

To identify potential targets of selection, we defined peak regions of amplification (regions of maximum recurrence e.g. **Fig. 2c,e>** top) within our CNAs and investigated the genes expressed in each of these regions. The eight peak regions contained between four and 397 expressed genes (FPKM > 0 in > 10% of lines) (**Fig. 2b,c,e**). We next filtered the list of putative targets using three criteria: (i) significant differential expression between lines with different copy numbers of the CNA region (ii) reported oncogenes from the COSMIC cancer gene census ^20^ or (iii) high scoring genes (in the top 2%) in a genome-wide siRNA of hES cell proliferation ^16^ (**Fig. 2b, c, e**). Using these criteria, we derived a candidate gene list that included *BCL2L1* on chr20, *EIF4A3*, BIRC5 (previously proposed as a putative target of selection by ^21^) and 10 others on chr17, and *DOCK1*, *SMNDC1* on chr10 (**Fig. 2b)**. Two genes on chr17, *EIF4A3* and *KPNB1*, scored more highly than *BCL2L1* in reducing proliferation after siRNA knock-down (top 0.1%) and were abundantly expressed in our iPSCs (top 98th and 99th mean expression percentile, respectively), but only one of these, *EIF4A3*, was found to be significantly over-expressed in lines with increased copy number (q = 7×10^−6^).

Duplications on chromosome 20 and 17 were also associated with changes in the expression of many genes outside of the CNA region, 80 and 984 respectively (false discovery rate 1%; FDR; **Fig. 2b,d**). Genes up regulated by the chr17 CNA were significantly enriched for members of the Notch signalling pathway. The Notch pathway may play a role in ESC proliferation ^22^ and down regulation of *NOTCH1* is associated with cell growth inhibition and increased apoptosis ^23^. Genes down regulated by the chr17 CNA were significantly enriched for genes involved in apoptosis modulation and signalling, including three members of the Bcl-2 protein family *BCL2L1* (pro/anti-apoptotic), BID (pro-apoptotic) and *PMAIP1/NOXA* (pro-apoptotic), the pro-apoptotic Bcl-2-interacting killer protein BIK, the pro-apoptotic genes *CASP9*, *DFFA* and *MAP3K5/ASK1* and the context-dependent pro/anti-apoptotic gene *DAXX* (FDR < 5%; **Supplementary Table 4**). Furthermore up-regulation of *EIF4A3*, the top target in the cis region of chr17, is thought to promote splicing of the anti-apoptotic BCL-XL isoform and the gene is known to regulate splicing of other apoptosis genes ^24^. In summary, we have produced the highest resolution map of recurrent CNAs in hiPSCs to date and identified a number of novel candidate genes that, when duplicated, may alter the growth properties of pluripotent stem cells, either by increasing proliferative capacity or decreasing apoptosis.

### Sources of hiPSC heterogeneity

We next explored how different technical and biological factors affect variation among iPSC lines using linear mixed models to partition the sources of variation of both Tier 1 and 2 assays (**Fig. 3a; Methods)**. Our experimental design included multiple independent lines from the same individual (136 donors with three lines, 37 more with two lines in Tier 1, 40 donors with two lines in Tier 2), enabling us to quantify between-individual differences (hereafter, ‘donor effects’) and to systematically compare this variance with that contributed by other factors. As expected, technical covariates, such as gene expression array batch, explained most variation in many of the assays. However, we also found consistent, statistically significant donor effects on the majority of iPSC phenotypes assayed, from methylation, through mRNA and protein abundances to cellular phenotypes such as pluripotency, differentiation capacity, and morphology (**Fig. 3b,c**). After accounting for technical batch factors, donor effects explained between 6.6% and 26.3% of the variance in the genome-wide assays averaging across all features in the assay, **Fig. 3a**), between 21.4% and 45.8% in the single protein immunostaining assays in pluripotent and differentiated cells(**Fig. 3b, S13**), and between 7.9% and 22.8% in the cellular morphology assays using an Operetta (Perkin Elmer) high content imaging system (**Fig. 3c**). Collectively, these results support the conclusion that differences between donor individuals affect the majority of iPSC cellular traits.

**Figure 3.**
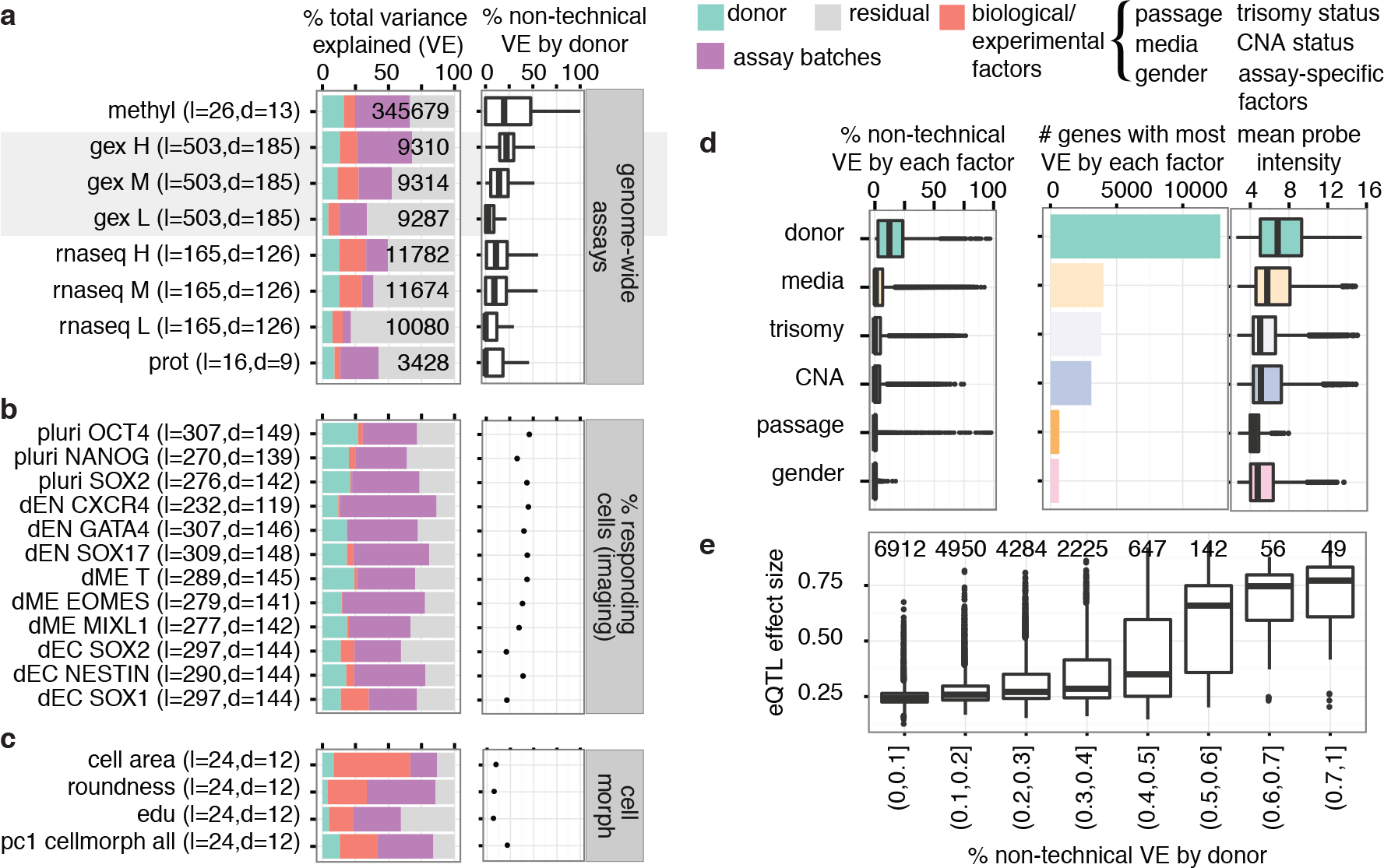
Variance component analysis of HipSci assays. **(a-c)** Variance component analysis for Tier-1 (270-522 lines) and Tier-2 assays (16-32 lines), partitioning phenotypic variability into donor effects, iPSC-specific experimental factors and assay batch. Left hand panels in **(a-c)** show the breakdown of total variance, right hand panels show proportion of variance explained by donor after accounting for technical covariates such as assay batches. For genomic assays, the average proportion of variance for genes in different abundance bins (medium, mid and high expressed)is shown. **(a)** genomic and proteomic assays **(b)** differentiation and pluripotency markers **(c)** cell morphology. **(d)** Breakdown of variance components of gene expression arrays from 522 Tier 1 lines excluding variance from assay batches. Left panel shows the distribution of the relative estimated donor, media, trisomy, CNA, passage number and gender variance components. Middle panel shows the number of genes where a particular variance component is the primary driver of heterogeneity (defined as the factor that explains to most variance). Right hand panel shows the mean gene expression level of the corresponding genes. **(e)** Relationship between donor variance component estimates and effect size estimates of lead eQTLs identified using gene expression arrays. Numbers above the boxplots denote the number of array probes in each variance bin.

We next investigated whether the variation we observed in iPSC gene expression could be further partitioned into additional biological factors after removing technical batch effects. Here we used data from Tier 1 gene expression arrays, the assay for which we have the largest number of donors and lines. Of the 25,391 remapped probes (17,011 genes) (**Methods; Supplementary Table 5**) measured, donor was the factor that explained the most variation in 51.8% of probes (52.1% of genes), substantially more than any other factor, including culture conditions (15.8%), trisomy status (15.1%), CNV status (12.2%), passage (2.6%) or gender (2.5%, **Fig. 3d**). Donor effects also appeared to be relatively consistent across all genes, while factors like culture condition or CNV status had large effects but only on a small numbers of genes (**Fig. 3d**). We observed only minor effects of gender on autosomal genes and line passage number explained little variation overall.

In principle, variation attributed to the donor in the variance component model maybe due to shared environment during reprogramming, in addition to common genetic background, because replicate lines were derived from the same population of fibroblast cells. To address this, we used the Tier 1 gene expression array data to map cis-acting expression quantitative trait loci (eQTL) in replicate lines from the same donor (**Supplementary Table 6**). We found that eQTL effect sizes were robust across replicate lines (**Extended Data Figure 5**), and large donor variation from the variance component model was associated with larger effect sizes of lead eQTL variants (**Fig. 3e, Methods**). This result strongly suggests that estimated donor variance components predominantly reflect genetic differences between donors.

### Identification of iPSC-specific regulatory variants

We next set out to characterise how the transcriptome of pluripotent cells is shaped by common genetic differences between individuals. We mapped expression quantitative trait loci (eQTL) using 167 iPSC lines from 127 unrelated donors using deep RNA-seq data, considering cis-acting variants within 1 Mb of the gene start. Genome-wide, we identified 2,169 genes with an eQTL at FDR 5% (hereafter referred to as ‘eGenes’) (**Supplementary Table 7; Methods**). Notably, power to discover eGenes in iPSCs was comparable to that in 44 somatic tissues studied by the GTEx Consortium ^25^ given our sample size (**Extended Data Figure 6; Supplementary Table 8**). Overall,iPSC eQTLs showed similar properties to eQTLs reported in other cell lines and somatic tissues (**Extended Data Figure 6**).

As many eQTLs are shared among tissues, we sought to place iPSC eQTLs in the broader context of somatic tissues. To define hiPSC-specific eQTLs we tested for replication of our eQTL signals in 44 tissues from GTEx, considering lead eQTL variants and their proxy variants (linkage disequilibrium r^2^ > 0.8; LD). Replication was defined using a nominal p < 0.01, Bonferroni adjusted for the total number of tissues tested. Using these criteria, we identified 503 eQTLs (503 eGenes) that were specific to iPSCs (**Fig. 4a; Methods**). We note that the proportion of iPSC-specific eQTLs (23%) was higher than in most GTEx tissues with comparable discovery sample size, with the exception of testis, a known outlier tissue ^26^. Notably, most of these signals (77%) occurred in genes with at least one reported GTEx eQTL that was notin high LD with the lead iPSC eQTL signal, suggesting that most iPSC specific eQTLs are driven by an alternative regulatory variant. Only 6% of the iPSC-specific eQTLs were explained by tissue-specific gene expression (**Methods**) (**Fig. 4b**), despite the known ubiquitous expression levels in iPS cells compared to somatic tissues (**Extended Data Figure 6**). Similar proportions were also seen when replicating eQTLs discovered in GTEx tissues. Interestingly, 20 of the iPSC-specific eQTLs regulate known cancer genes (Fisher p = 6.7×10^−4^ compared to eGenes regulated by non-specific eQTLs), including the tumour-suppressor *TP53* (**Supplementary Table 9**). For three of these genes (*NRAS*, *HNF1A*, and *NFATC2*) there was no eQTL detected in any other GTEx tissue. Compared to tissue-specific effects in GTEx tissues, iPSC-specific eQTLs appeared to regulate more cancer-implicated genes than somatic tissues (**Extended Data Figure 7**). We also found an iPSC-specific eQTL with a large effect size for *BIRC5* (**Extended Data Figure 7**), a gene that is commonly overexpressed in tumours and identified as one of the candidate genes under selection by a recurrent CNA on chromosome 17 (**Fig. 2e**).

For a subset of iPSC-specific eGenes we observed a corresponding effect on protein abundance, although the small number of lines with proteomics data (**Extended Data Figure 3**) prevented genome-wide analysis of proteome quantitative trait loci (pQTL). An example is shown for rs10999085 targeting the *H2AFY2* (*H2A Histone Family, Member Y2*) gene (**Fig. 5a,b**), which encodes for a replication-independent histone protein that functions in transcriptional repression and has been connected with differentiation ability in pluripotent cells^27^. Taken together, our results suggest that gene regulation in iPSCs is partly driven by iPSC-specific regulatory elements, in line with a recent study assessing self-renewal capacity in ESCs and macrophages ^28^.

**Figure 4.**
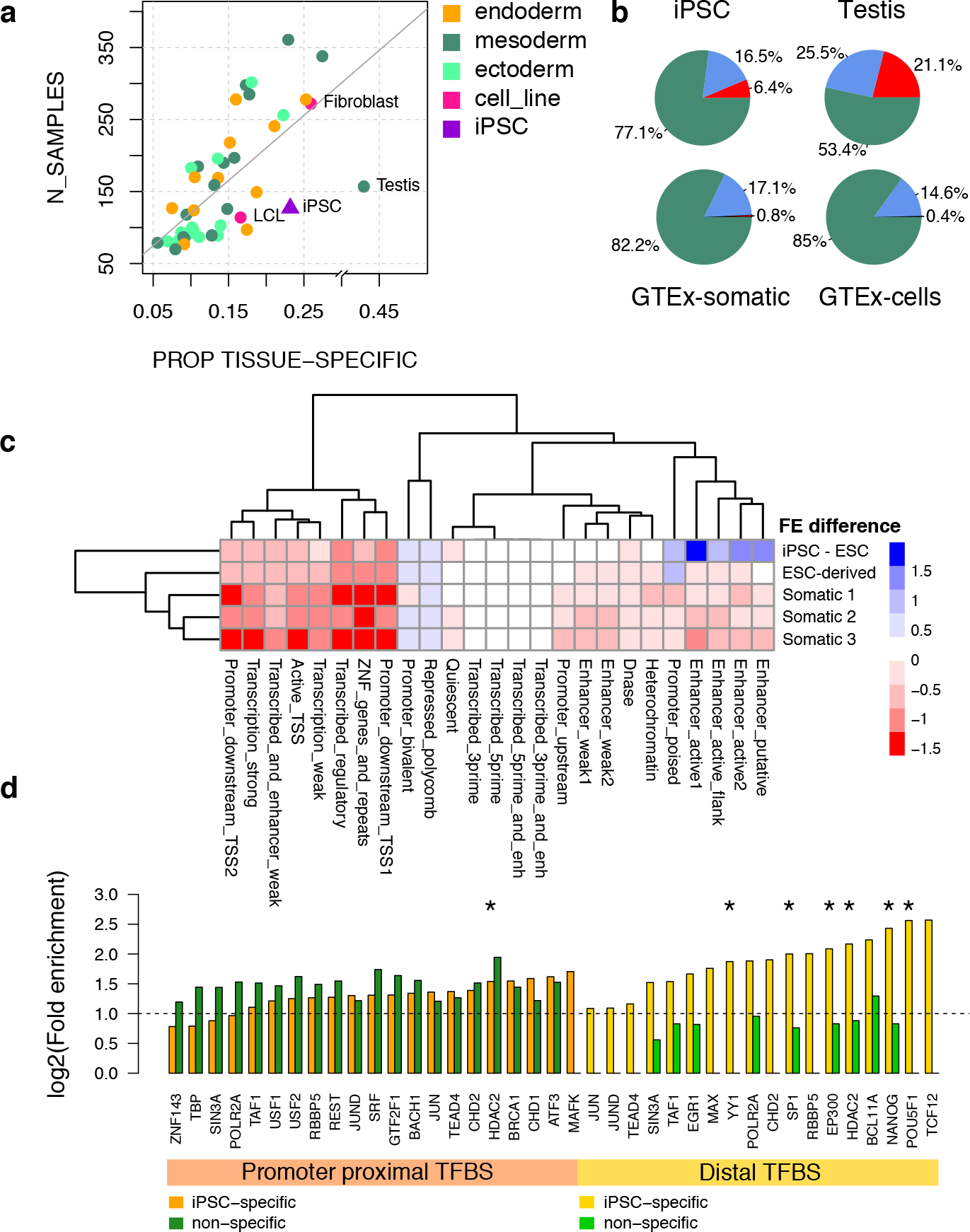
iPSC eQTL map in the context of somatic tissues. **(a)** Proportion of tissue-specific eQTLs (considering the replication of lead eQTLs and their high-LD proxies; r^2^ > 0.8) as a function of the discovery sample size. Points other than iPSC (this study) are from the GTEx Consortium (44 somatic tissues and cell lines) ^25^. **(b)** Assignment of the most likely causes for tissue-specific eQTLs shown for iPSCs, GTEx testis and the average of GTEx somatic tissues and cell lines. Breakdown: gene not expressed (red); gene expressed but not no eQTL (blue); eQTL effect is driven by distinct lead variants (r^2^ < 0.8; green). **(c)** Heatmap of the fold enrichment (FE) difference between iPSC-specific and non-specific eQTLs at 25 chromatin states from the Roadmap Epigenomics Project ^29^, shown for five aggregated clusters representing 127 different cell types. Colouring: higher FE in iPSC-specific (blue), higher FE in somatic (red). **(d)** Enrichment of iPSC eQTLs at promoter proximal and distal (defined as less than or greater than 2 kb from the transcription start site) transcription factor binding sites (TFBS) in H1-hES cells from the ENCODE Project ^30^. Significant fold enrichments per factor are shown for iPSC-specific and non-specific eQTLs. Pluripotency-associated factors are indicated with an asterisk.
Functional genomic context of iPSC-specific eQTLs

### Functional genomic context of iPSC-specific eQTLs

The transcriptional regulatory networks that maintain pluripotency are unique to stem cells. We next investigated how common variants modulate these networks to produce iPSC-specific genetic effects on expression. We used chromatin state annotations from 127 reference epigenomes from the Roadmap Epigenomics Project ^29^ to quantify the fold enrichment of iPSC-specific and nonspecific eQTL sets across all chromatin states relative to randomized matched variants (**Methods**). iPSC-specific eQTLs were highly enriched in two clusters: active enhancers and poised promoters in pluripotent stem cells and in ESC-derived cell types, primarily the three embryonic germ layers. In contrast, eQTLs that are not tissue-specific were most highly enriched near active promoters and transcribed regions across different somatic tissues (**Fig. 4c**). iPSC-specific eQTLs were also significantly enriched for binding sites of key regulators of pluripotency obtained from the ENCODE Project ^30^, including *NANOG, POU5F1* (*OCT4*), and multiple other factors relevant for pluripotency ^9, 31^, compared with non-specific eQTLs which did not show comparable enrichment for these factors (**Methods; Fig. 4d**). This enrichment was predominantly seen at distal transcription factor binding sites (defined as > 2 kb away from the TSS), in accordance with previous observations of tissue and context-specific regulatory elements being more likely distal than proximal ^32, 33^. Our results suggest that common genetic differences between individuals may affect regulation during early stages of development.

### iPSC eQTLs tag common disease variants

Although the value of iPSCs for genetic engineering experiments is clear, much less is known about their relevance as a model cell type for functional interpretation of common disease-associated variants. To explore this, we overlapped all iPSC eQTLs (lead variants and their high-LD proxies) with the NHGRI-EBI catalogue for genome-wide association studies (GWAS). iPSC eQTLs and their proxies tagged a total of 85 catalogued variants associated with 67 different traits. Amongst the 85 variants there were 46 distinct loci for which the eQTL effect was strongest in iPSC cells, and 8 loci that were tagged by iPSC-specific eQTLs (**Supplementary Table 10; Methods**). Globally, this numberof tagging events was similar to what is expected by chance (using randomized eQTL variants matched for allele frequency, distance to the nearest transcription start site, gene density, and number of LD proxy variants; **Methods**). However, when considering individual traits, we found eQTLs to be enriched for variants associated with12 traits (minimum two variants; **Supplementary Table 10**).

**Figure 5.**
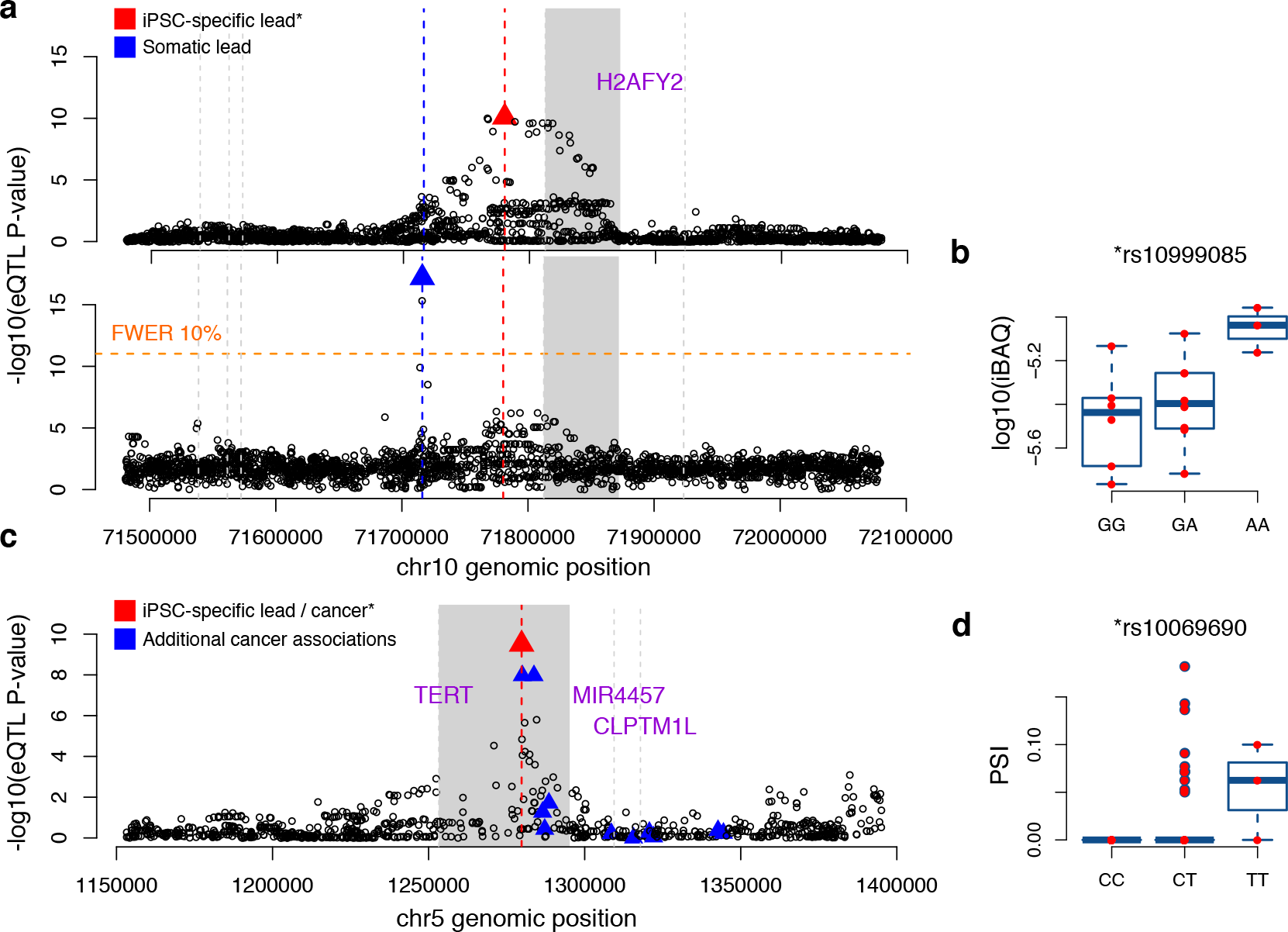
iPSC-specific eQTLs tag disease-associated variation. **(a)** Example of an iPSC-specific eQTL locus, highlighting the lead eQTL variant rs10999085 (red), the target eGene (*H2AFY2*; H2A Histone Family, Member Y2; gray; upper panel), and somatic eQTL signal at the same locus (lower panel). Shown is the-Iog10(minimum eQTL p-value) derived from 44 GTEx tissues, highlighting the distinct loci driving the regulation of *H2AFY2* in iPSCs and somatic tissues. The orange horizontal line indicators the family-wise error rate (FWER) of 10% (**Methods**). Start positions for other protein-coding genes are indicated with vertical grey lines, **(b)** Replication of the *H2AFY2* eQTL on the protein level, showing the log 10 scaled iBAQ values for 17 iPSC lines (9 donors) (pQTL nominal p = 0.0085; linear regression), stratified by their genotype at rs10999085. **(c)** Example of a disease-tagging iPS specific eQTL locus on chromosome 5. The disease variant rs10069690 is associated with multiple different types of cancer, including breast and ovarian cancer ^34,35^ and tagged by an eQTL for *TERT* (Telomerase Reverse Transcriptase). The lead eQTL variant is highlighted in red and additional cancer-associated variants in blue. The gene region of *TERT* in indicated in solid gray and transcription start sites for other protein-coding genes in the region are shown with verticalgray lines, **(d)** Boxplot showing *TERT* intron 4 retention ratio (PSI, percent spliced in) in iPSC lines of all individual donors stratified by their genotype at rs10069690.

We conclude with one example of a GWAS variant that shows an iPSC-specific eQTL effect, which illustrates how studying the genetic regulation of gene expression in iPSCs may help generate insights into the mechanisms through which GWAS disease variants act. Variant rs10096960 on chromosome five is a lead eQTL variant for the *TERT* (*Telomerase Reverse Transcriptase*) gene, which encodes the catalytic subunit of the human telomerase enzyme (**Fig. 5c, d**). This variant is associated with germline predisposition to seven different cancers ^34– 36^, and there are multiple additional variants at the same locus associated with cancer as well as other phenotypes such as telomere length ^37,38^. *TERT* promoter mutations are also the most frequent non-coding somatic mutations observed in a variety of cancers such as melanoma ^39^. To explore putative mechanistic effects of rs10069690, we analysed alternative splicing of the *TERT* gene, as previous studies have reported aberrant splicing caused by this variant ^38^ as well as highly abundant alternative *TERT* transcripts in ESCs ^40^. We quantified *TERT* intron retention rates and identified two alternative splicing events associated with rs10069690 (i.e. splicing QTLs). One of them affects the intron where the variant is located, with the minor allele of rs10069690 (T), increasing the fraction of *TERT* transcripts in which intron 4 is retained (p = 4.6e-05, Bonferroni adjusted) (**Methods; Fig. 5d, Extended Data Figure 8**). Recent work has shown that an increase in *TERT* expression caused by regulatory promoter mutations only manifests in differentiated cells, where increased *TERT* expression results also in increased telomerase activity ^41^. We therefore speculate that the eQTL affecting *TERT* expression in iPSCs results in genotype-dependent variability in telomerase activity in somatic cell types, possibly mediated by aberrant splicing, which leads to differential cancer susceptibility.

## Discussion

Here we present the first analyses of genetic and phenotypic data from 522 human iPSC lines derived by HipSci. Our study illustrates that iPSC technology is sufficiently mature to generate high quality cell lines from hundreds of individuals, facilitating large-scale studies of the consequences of human genetic variation in pluripotent stemcells. Strikingly, our data demonstrate that donor effects are a major driver of molecular and cellular heterogeneity in iPSCs after accounting for technical batch effects.While interindividual variation in gene expression is perhaps not surprising, our data suggest that genetic differences affect a wide range of molecular and cellular phenotypes, including the efficiency at which iPS cells differentiate into the three embryonic germ layers ^42– 44^. One interpretation of this finding is that common genetic variation has subtle effects on core components of the regulatory networks controlling cellular differentiation and responses to external environmental stimuli. A major advantage of genetic studies in iPSCs compared to other immortalised cell lines such as EBV-transformed lymphoblastoid cell lines ^45– 47^ is that effects can be analysed and compared in different derived cell types, while sharing genetic data. Future efforts to map quantitative trait loci that regulate these networks will provide a novel and powerful tool for dissecting the genetic architecture of development and somatic tissue physiology.

We have generated the most extensive map so far of the locations of recurrent genetic abnormalities in iPSCs. Compared to previous large-scale studies in human embryonic and induced pluripotent stem cells ^6, 11^, we observed lower levels of genetic aberrations in our lines. One possibility for this difference is that previous studies have primarily focused on cells with relatively high passage numbers compared to our hiPSCs,although we did not notice a significant increase in rate of CNAs with passage number in our study. Alternatively, due to the lack of reference donor samples in previous work, some germline CNAs might have been mistaken for events that occurred during reprogramming and cell culture. Indeed, even within our study, the CNAs could have occurred somatically in the donor prior to skin biopsy, and been selected either in the donor or in the reprogramming and cell growth process, as has been suggested by recent work ^12^. This would be consistent with the same variant appearing in separately selected iPSC lines from the same donor fibroblast pool. We were also able to link a subset of the CNAs to downstream transcriptional changes, the most notable association being between CNAs on chromosome 17 and changes in genes that regulate apoptosis and Notch signalling, suggesting that these changes may result from a growth advantage conferred by duplication of specific genes.

Our study provides the first map of common regulatory variation in human pluripotent cells. We show that variation in local gene regulation in iPSCs is similar to that in somatic tissues, with eQTLs driving lineage-specific expression profiles through distal tissue-specific regulatory elements such as enhancers. We have identified a set of variants that show regulatory function only in pluripotent cells and identified complex disease-associated loci tagged by these variants. These loci may drive disease-susceptibility through molecular changes early in development or more generally in cells with ‘stem-like’ characteristics, which are not well captured by studies of differentiated primary tissues from adult individuals. A compelling example of this is the iPSC-specific eQTL regulating *TERT* expression. In human tissues, telomerase activity is mainly restricted to stem cells, with most somatic tissues silencing *TERT* expression. However, cancer cells bypass this tumor suppressive mechanism by reactivating telomerase activity ^48^, returning toa more ‘stem-like’ state. This result highlights how iPSCs could be used to study the genetic effects of diseases that manifest in transient states during cellular growth and differentiation, which are known to be particularly important in cancer^49^

In summary, our study provides a first comprehensive picture of the genetic and phenotypic variability in human pluripotent stem cells, including the major drivers of this variation. Data and cell lines from this study are being made available through www.hipsci.org and the European Collection of Authenticated Cell Cultures (ECACC). As the HipSci resource continues to expand in sample size and assays, it will enable the study of more subtle genetic effects, under a wider range of conditions, in an increasing range of disease-relevant differentiated cell types.

## Methods

All samples for the HipSci resource were collected from consented research volunteers recruited from the NIHR Cambridge BioResource (http://www.cambridgebioresource.org.uk). Samples were collected initially under existing Cambridge BioResource ethics for iPSC derivation (REC Ref: 09/H0304/77, V2 04/01/2013), with later samples collected under a revised consent (REC Ref: 09/H0304/77, V3 15/03/2013). Details of the generation and phenotyping of all cell lines used in the study, data generation and analysis are described in the Supplementary Information.

## Acknowledgements

This work was funded with a strategic award from the Wellcome Trust and Medical Research Council (WT098503). We thank the staff in the Cellular Genetics and Phenotyping and Sequencing core facilities at the Wellcome Trust Sanger Institute. Work at the Wellcome Trust Sanger Institute was further supported by Wellcome Trust grant WT090851. FMW gratefully acknowledges financial support from the Department of Health via the NIHR Biomedical Research Centre award to Guy's & St Thomas' National Health Service Foundation Trust in partnership with King's College London and King's College Hospital NHS Foundation Trust. We gratefully acknowledge the participation of all NIHR Cambridge BioResource volunteers, and thank the NIHR Cambridge BioResource centre staff for their contribution. We thank the National Institute for Health Research and NHS Blood and Transplant. The volunteer recruitment was supported by the NIHR/Wellcome Trust Cambridge Clinical Research Facility.

### Author contributions

HK, AG, OS, DG: Wrote the paper with input from all authors.

HK, AG, DB, YM, IS, PD, DMcC, AA, MP, DD, AL, OS, DG: Contributed to the supplementary material

HK, AG, AL, FPC, PD, DMcC, DD: Analysed the data

SA, WO: Managed and supervised collection of research volunteer samples FS, CA, AA, RN, SH, MP, CK: Generated iPSC lines, Tier 1 assay data, RNA-seq and methylation data

VA, DB: Generated and processed the proteomics data

AL, OC, RM, NM, DD: Generated and processed the high content cellular imaging data

SMcC, YM: Initial data quality control and bioinformatics processing/pipelines AF, PH, IS, LC: Curated and managed data and project website RH, AKK: Coordinated the project

DG, OS, RD, FW, AIL, LV, EB, WO, PB, DD: Supervised and designed the research

### Author information

Details of the data generated during the project, including archive accession identifiers for obtaining the data, are described in the Supplementary Information. The HipSci website (www.hipsci.org) also has full details of all publicly available data and instructions for researchers to register for access to data in European Genome-phenome Archive (EGA). The authors declare no competing financial interests. Correspondence and requests for materials should be addressed to.

